# Pseudotyped Bat Coronavirus RaTG13 is efficiently neutralised by convalescent sera from SARS-CoV-2 infected Patients

**DOI:** 10.1101/2021.08.17.456606

**Authors:** Diego Cantoni, Martin Mayora-Neto, Nazia Thakur, Ahmed ME Elrefaey, Joseph Newman, Sneha Vishwanath, Angalee Nadesalingam, Andrew Chan, Peter Smith, Javier Castillo-Olivares, Helen Baxendale, Bryan Charleston, Jonathan Heeney, Dalan Bailey, Nigel Temperton

## Abstract

RaTG13 is a close relative of SARS-CoV-2, the virus responsible for the Coronavirus Disease 2019 (COVID-19) pandemic, sharing 96% sequence similarity at the genome-wide level. The spike receptor binding domain (RBD) of RaTG13 contains a large number of amino acid substitutions when compared to SARS-CoV-2, likely impacting affinity for the ACE2 receptor. Antigenic differences between the viruses are less well understood, especially whether RaTG13 spike can be efficiently neutralised by antibodies generated from infection with, or vaccination against, SARS-CoV-2. Using RaTG13 and SARS-CoV-2 pseudotypes we compared neutralisation using convalescent sera from previously infected patients as well as vaccinated healthcare workers. Surprisingly, our results revealed that RaTG13 was more efficiently neutralised than SARS-CoV-2. In addition, neutralisation assays using spike chimeras and mutants harbouring single amino acid substitutions within the RBD demonstrated that both spike proteins can tolerate multiple changes without dramatically reducing how efficiently they are neutralised. Moreover, introducing the 484K mutation into RaTG13 resulted in increased neutralisation, in contrast to the same mutation in SARS-CoV-2 (E484K). This is despite E484K having a well-documented role in immune evasion in variants of concern (VOC) such as B.1.351 (Beta). These results indicate that the immune-escape mutations found in SARS-CoV-2 VOCs might be driven by strong antibody pressures, and that the future spill-over of RaTG13 and/or related sarbecoviruses could be mitigated using current SARS-CoV-2-based vaccination strategies.

## Introduction

The Coronavirus disease 2019 (COVID-19) pandemic, caused by the severe acute respiratory syndrome coronavirus 2 (SARS-CoV-2), has currently surpassed 150 million recorded cases, claimed upwards of 3 million lives and continues to overwhelm health-care facilities in countries around the world (1,2). SARS-CoV-2 is a betacoronavirus, as are the close relative SARS-CoV and Middle Eastern respiratory syndrome-related virus (MERS), both of which have spilled over into human populations from animal reservoirs. The natural reservoir of alpha and betacoronaviruses is widely believed to be bats. The direct ancestor of SARS-CoV-2 remains to be identified, as well as a possible intermediate reservoir. More than 250 coronaviruses have been detected in bats (3).

In 2013, a bat coronavirus was detected in Mojiang County, Yunnan, China, eventually denoted as RaTG13. Following the emergence of SARS-CoV-2 in 2019, the RaTG13 isolate was shown to be one of the closest known relatives, with 96.2% sequence similarity at the genome-wide level (1). Of note, isolates of related viruses in pangolins have receptor binding domains (RBDs) which are more closely related, likely a broader reflection of frequent sarbecovirus recombination in reservoirs (4). To date, the direct ancestor of SARS-CoV-2 remains to be identified, as well as any possible intermediate reservoir. Nevertheless, SARS-CoV-2 and RaTG13 both rely on their Spike protein for viral entry into cells, via the cell surface angiotensin-converting enzyme 2 (ACE2) receptor, complemented by activity of additional proteases including Transmembrane protease, serine 2 (TMPRSS2). Comparative studies of the structures of both SARS-CoV-2 and RaTG13 Spike revealed a high degree of conservation in the ectodomain (97.8%) with most of the substitutions being within the RBD, clustering around the ACE2-binding interface and the RBD-RBD interfaces of the trimeric Spike complex (5,6). Previously, we have identified that introducing SARS-CoV-2 changes into RaTG13 Spike increases the tropism for human ACE2, providing mechanistic insight into the potential pathway for spill-over (7). Of note, these mutants were constructed in Spike expression constructs for pseudotyping and were not inserted into recombinant RaTG13 or SARS-CoV-2 viruses. Given the structural similarities between SARS-CoV-2 and RaTG13 Spike, as well as the likelihood of a shared ancestor, we wanted to assess whether antibodies generated following prior infection with SARS-CoV-2 or through Spike-protein based vaccination, would result in neutralisation of RaTG13. This is particularly relevant in the context of emerging variants of concern (VOCs), e.g. B.1.1.7 (Alpha), B.1.351 (Beta) and B.1.617.2 (Delta) as many of the amino acid substitutions between RaTG13 and SARS-CoV-2 Spike are at positions that have been established as important (antigenically or functionally) in VOCs, e.g. 484 and 501. It is essential we develop an understanding of the breadth of immunity conferred by both infection and vaccination. Indeed, knowledge on whether closely related coronaviruses to SARS-CoV-2 are neutralised by current vaccination programs can be used to gauge the risk of potential sarbecovirus spillover events in the future.

To characterise the degree of cross-neutralisation between these related Spikes, we pseudotyped SARS-CoV-2 and RaTG13 using a lentiviral core and performed pseudovirus microneutralisation assays (pMN) to assess the neutralisation potency of convalescent sera derived from previously infected SARS-CoV-2 patients and vaccinated healthcare workers (HCWs) (sampled in the United Kingdom). Surprisingly, despite the significant number of RBD substitutions in RaTG13 these pseudotypes were more efficiently neutralised by our sera. These results have important implications when anticipating further zoonotic transmission of coronaviruses and the likely protection afforded by current vaccination approaches and immune responses to SARS-CoV-2.

## Materials and Methods

### Tissue Culture

Human Embryonic Kidney 293T/17 (HEK293T/17) cells were cultured using Dulbecco’s modified eagle medium (DMEM, PanBiotech) supplemented with 10% foetal bovine serum (PanBiotech) and 1% penicillin/streptomycin (PanBiotech), in a 37°C, 5% CO_2_ incubator. Cells were routinely passaged three times a week to prevent overconfluency.

### Plasmid generation

The RaTG13 construct, the chimeric expression plasmids expressing SARS-CoV-2 with the RaTG13 RBD and RaTG13 with the SARS-CoV-2 RBD as well as the individual mutants were synthesised commercially (BioBasic) and subcloned into pcDNA3.1+ expression vectors, as detailed in Conceicao *et al*, 2020 (7). The B.1.351 (Beta) variant Spike was synthesised commercially (GeneArt) and subcloned into a pCAGGS expression vector. The SARS-CoV-2 Spike expression plasmid was kindly gifted by Professor Xiao-Ning Xu, Imperial College, London.

### Pseudotype virus generation

The generation of lentiviral pseudotyped viruses (PV) bearing the Spike protein of either SARS-CoV-2 or RaTG13 was adapted from a previously described protocol (8). The day prior to generating PVs, cells were seeded in T-75 flasks for next day transfection at a density of 70% confluency. On the day of transfection, a DNA mix containing 1000ng of p.891 HIV Gag-Pol, 1500ng of pCSFLW luciferase and 1000ng of either SARS-CoV-2 Spike, RaTG13 Spike, Spike chimeras or B.1.351 (Beta) variant were prepared in 200µL of OptiMEM, followed by addition of transfection reagent FuGENE HD (Promega) at a 1:3 ratio and incubated for 15 minutes. During this time, culture media was replenished, and transfection complexes were added dropwise into the culture flasks. PVs were harvested 48 hours post transfection by filtering culture media through a 0.45µm cellulose acetate filter. PVs were then titrated and aliquoted for storage at -70°C.

### Pseudotype virus titration

Titration of PVs was carried out as previously described (8). Target cells were prepared the day prior to titration of PVs by transfecting HEK203T/17 cells with 2000ng ACE2 and 150ng of TMPRSS2 plasmids using FuGENE HD, to render cells permissible to SARS-CoV-2 and RaTG13 PVs. To titrate PVs, 100µL of undiluted PVs were serially diluted 2-fold down a white 96-well F-bottom plates (Perkin Elmer) in 50µL of DMEM. Target cells were added at a density of 10,000 cells were per well, and plates were returned to the incubator for 48 hours prior to lysis using Bright-Glo reagent (Promega). Luminescence was measured using a GloMax luminometer (Promega). PVs were quantified based on relative luminescence units per ml (RLU/ml).

### Neutralisation Assays

pMNs were carried out as previously described (8). Briefly, heat inactivated human convalescent sera was diluted 1:40 with DMEM and serially diluted 2 fold down white 96-well F-bottom plates. PVs were added at a density of 5×10^5^ RLU/ml in each well and incubated for 1 hour at 37°C and 5% CO_2_, prior to the addition of target cells at a density of 10,000 cells per well. Plates were returned to the incubator for 48 hours prior to lysis with Bright-Glo reagent. Luminescence was measured using a GloMax luminometer (Promega). Data was analysed using GraphPad Prism software to derive IC_50_ values, as previously described (9).

### Serum Sample Collection

Serum samples were obtained from healthcare workers (HCW) working at Royal Papworth Hospital, Cambridge, UK (RPH) and from COVID-19 patients referred to RPH for critical care. (Study approved by Research Ethics Committee Wales, IRAS: 96194 12/WA/0148. Amendment 5). HCWs from RPH were recruited through staff email over the course of 2 months (20th April 2020-10th June 2020) as part of a prospective study to establish seroprevalence and immune correlates of protective immunity to SARS-CoV-2. Patients were recruited in convalescence either pre-discharge or at the first post-discharge clinical review. Patient sera (n=20) and seropositive HCW sera (n=5) were obtained between 6^th^ of June 2020 and 22^nd^ of September 2020. Sera samples from HCW immunised with single (n=21) and double doses (n=20) of either Pfizer BNT162b2 (1^st^ dose: n=12, 2^nd^ dose: n=12) or AZD1222 (1^st^ dose: n=9, 2^nd^ dose: n=11) vaccines were obtained 4 to 6 weeks after each dose of vaccination. All participants provided written, informed consent prior to enrolment in the study.

At recruitment, HCW were classified as pre-exposed according to the results provided by a CE-validated Luminex assay detecting N-, RBD- and S-specific IgG, a lateral flow diagnostic test (IgG/IgM) and an Electro-chemiluminescence assay (ECLIA) detecting N- and S-specific IgG as previously described in (10). Any sample that produced a positive result by any of these assays was classified as positive.

## Results

To assess differences in neutralisation between SARS-CoV-2 and RaTG13, we initially used the WHO International Reference Panel for anti-SARS-CoV-2 immunoglobulin (NIBSC; 20/268) which provides five sets of sera, one negative, and four positive, categorised by increasing amounts of neutralising antibodies specific to SARS-CoV-2. Interestingly, we observed higher neutralising potency against RaTG13, when compared to SARS-CoV-2 in three of the four sera samples, not including the negative sera control (Figure 1A). Switching to a larger cohort and repeating the assay with 25 convalescent sera samples derived from patients and healthcare workers (HCWs) who were infected with SARS-CoV-2 during the first wave in the United Kingdom, we observed a similar trend, with RaTG13 being more efficiently neutralised when compared to SARS-CoV-2 (2.2 fold change, p=<0.0001) (Figure 1B). In comparison, B.1.351 (Beta), a VOC thought to have emerged in South Africa, showed a significant reduction in neutralisation compared to SARS-CoV-2 (6.4 fold change, p=<0.0001), confirming previous observations (11–13).

**Figure 1.**
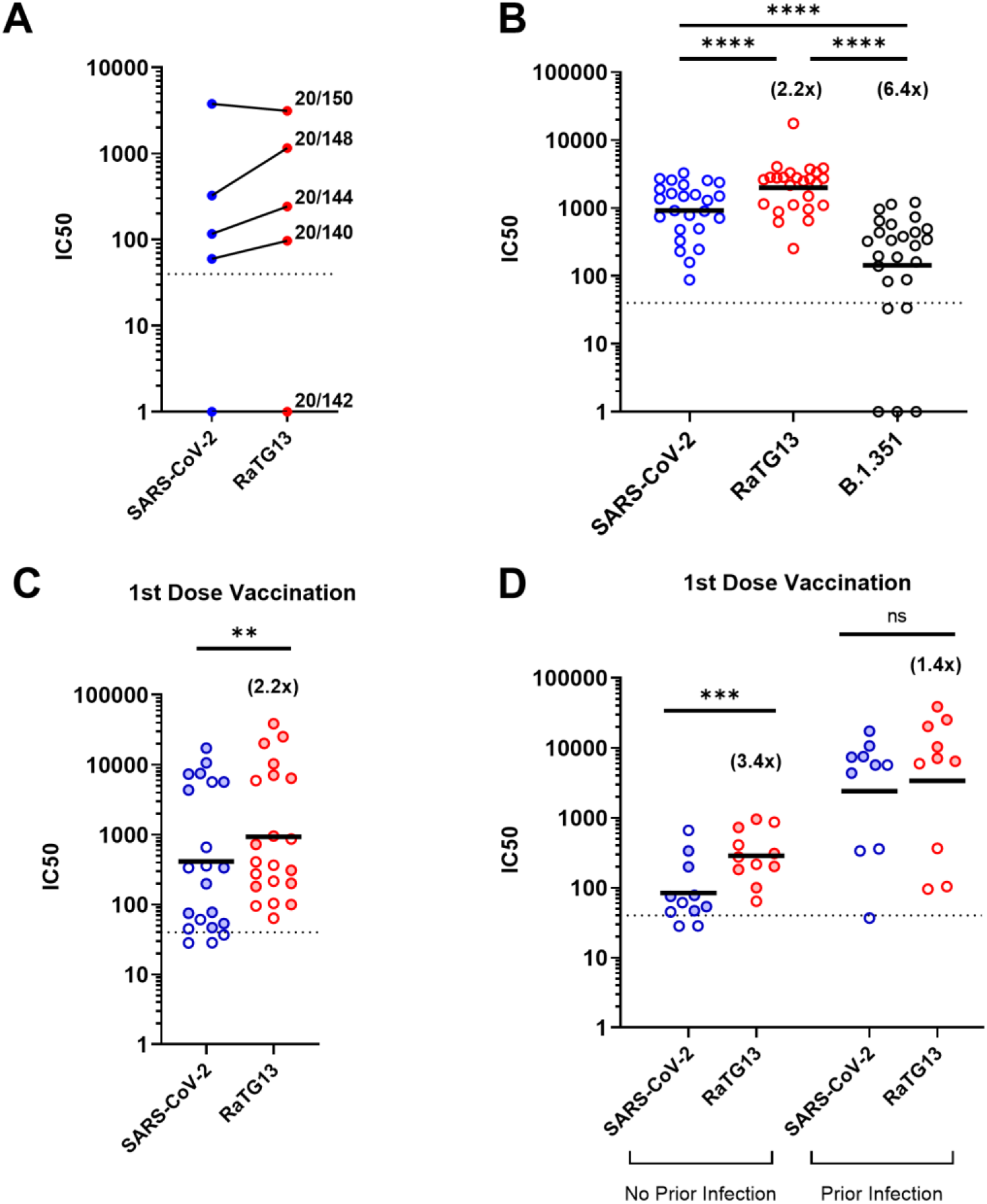
Differences in neutralisation titres between SARS-CoV-2 and RaTG13 by pMN Assay. (A) Comparison of neutralisation titres between SARS-CoV-2 and RaTG13 using the WHO International Reference Panel for anti-SARS-CoV-2 immunoglobulin. Three of the four sera showed increased neutralisation titres against RaTG13. (B) Comparison of neutralisation titres between SARS-CoV-2, RaTG13 and the B.1.351 (Beta) variant of concern using convalescent sera derived from patients and healthcare workers. (C) Comparison of neutralisation titres between SARS-CoV-2 and RaTG13 against sera from single dose vaccinated healthcare workers. (D) Differences in neutralisation titre from single dose vaccinated healthcare workers split by ‘no prior infection’ or ‘prior infection’ with SARS-CoV-2. Full circles denote healthcare workers vaccinated with BNT162b2, whereas open circles denote vaccination with AZD1222. Numbers in brackets denote fold changes relative to SARS-CoV-2. Wilcoxon matched pairs signed rank tests were used in panels B, C and D. Dotted lines in graphs denote the assay’s lower limit of detection. IC50 was calculated by fitting a non-linear regression curve using Graphpad Prism 8 software.

To compare these results to vaccine-derived immunity, we examined sera from HCWs who had received a first dose of either the BNT162b2 (Pfizer-BioNTech, n=12) or AZD1222 (Oxford-AstraZeneca, n=9) vaccine against SARS-CoV-2. pMN assays were carried out on post-vaccination sera samples (n=21), which again revealed higher neutralisation titres against RaTG13 compared to SARS-CoV-2 (2.2 fold change, p=0.0016) (Figure 1C). This difference was recapitulated when stratifying the group based on the absence of prior infection (n=11) (3.4 fold change, p=0.001) (Figure 1D). Interestingly, the difference in geometric IC_50_ means between SARS-CoV-2 and RaTG13 in HCWs with documented prior infection (n=10) was not significant (1.4 fold change, p=0.084) (Figure 1D). Together these data provide compelling evidence that vaccination or natural infection provides cross-protective immunity to RaTG13, at least at the level of neutralising antibodies.

To examine the differences in viral neutralisation in more detail we utilised two SARS-CoV-2/RaTG13 chimeric Spike plasmids (7). These two chimeric Spikes (SARS-CoV-2 Chimera: N439K, Y449F, E484T, F486L, Q493Y, Q498Y, N501D, Y505H, and RaTG13 Chimera: K439N, F449Y, T484E, L486F, Y493Q, Y498Q, D501N, H505Y) contain the amino acid substitutions between the two viruses that are present within the RBD and known to interact with human ACE2 (5,14) (Figure 2A, B). A number of these residues are implicated in the evasion of neutralisation, either with the same amino acid change, e.g., N439K (15) or different, e.g., N501Y and E484K are seen in B.1.351 (16), as opposed to N501D and E484T found in RaTG13. Analysing the same patient and HCW convalescent sera set (n=25; Figure 1), we identified that the SARS-CoV-2 Chimera was neutralised more efficiently than SARS-CoV-2 WT (1.6 fold change, p=0.0005). In contrast, the RaTG13 Chimera was neutralised slightly less efficiently than RaTG13 WT (1.2 fold change, p=0.0043) (Figure 2C). These relative changes in neutralisation efficiency indicated that differences in neutralisation between SARS-CoV-2 and RaTG13 (WTs) might therefore be attributable, in part, to amino acid substitutions within the chimeras.

**Figure 2.**
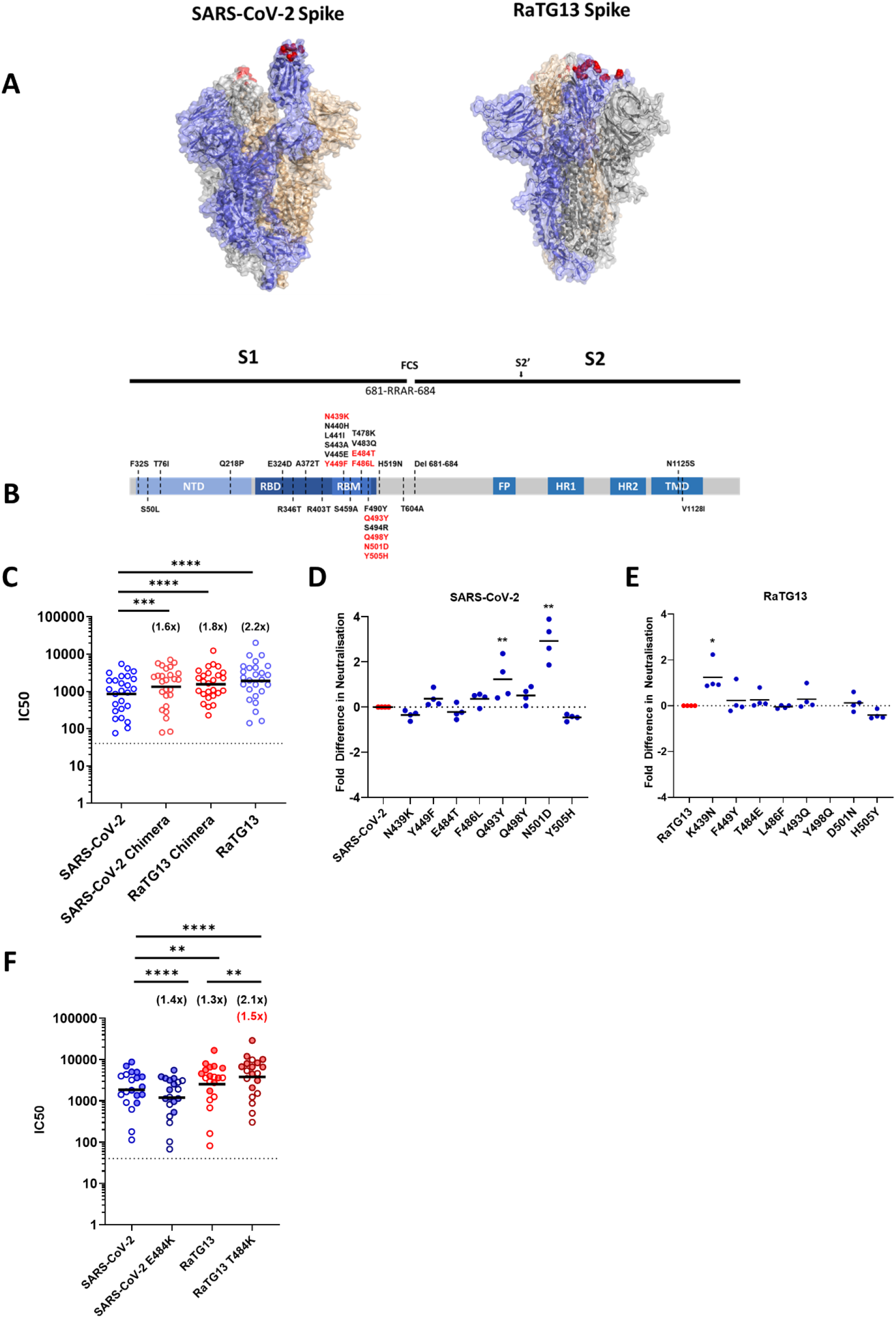
Key amino acid residues affect antibody neutralisation against SARS-CoV-2 and RaTG13. (A) Structures of SARS-CoV-2 (PDB: 6X2A) and RaTG13 (PDB: 7CN4). The highlighted (red) amino acids denote the ACE2 contact residues that were substituted to generate the chimera plasmids for subsequent pMN assays. (B) Simplified schematic highlighting the functional domains within the SARS-CoV-2 Spike protein. The amino acid substitutions found in RaTG13 have been labelled across the diagram, with red text denoting the substitutions displayed in Panel A, which were built into chimeric Spikes. (FCS, furin cleavage site; NTD, N-terminal domain; RBD, receptor binding domain; RBM, receptor binding motif; FP, fusion peptide; HR, heptad repeat; TM, transmembrane domain). (C) Repeated experiments showing the neutralisation titres, by pMN assay, of SARS-CoV-2, RaTG13 and both chimeras with the same set of convalescent sera used in Figure 1. (D, E) Using a set of four convalescent sera derived from patients, each pseudotype mutant was assayed by pMN and IC50s were then converted to fold changes against their original SARS-CoV-2 (D) or RaTG13 (E) background. RaTG13 Y498Q was not done due to low PV titre. (F) Lysine substitution at position 484 does not decrease neutralisation of RaTG13, but does so for SARS-CoV-2. Comparisons of neutralisation were made using sera from doubly vaccinated HCWs. Full circles denote healthcare workers vaccinated with BNT162b2, whereas open circles denote vaccination with AZD1222. Numbers in brackets denote fold changes relative to SARS-CoV-2, with the exception of the red coloured values which denote fold change between RaTG13 and RaTG13 T484K. Wilcoxon matched pairs signed rank tests was used in panel C. A Student’s test was used in panels D and E. IC50 was calculated by fitting a non-linear regression curve using Graphpad Prism 8 software.

To identify the specific changes responsible for RaTG13’s enhanced neutralisation we next examined each substitution in isolation (Figure 2D and E), comparing their neutralisation to WT virus with four randomly chosen patient sera samples. For RaTG13 we found that the substitution of individual amino acids had little appreciable effect on neutralisation, with the exception of K439N, which resulted in a mean 1.2 fold increase in neutralisation (p=0.0194) (Figure 2D). In contrast, several individual substitutions in the SARS-CoV-2 background (Figure 2E) showed significant increased neutralisation potency, with Q493Y showing a mean 1.2 fold increase (p=0.0088) and N501D a mean 2.9 fold increase (p=0.0069). These results highlight the importance of these particular residues in defining the relative neutralisation of RaTG13 and SARS-CoV-2 by convalescent sera. RaTG13 Y498Q neutralisations were not performed due to insufficient viral titres being attained during pseudotype preparation.

Lastly, the established importance of SARS-CoV-2 E484K substitutions in immune-escape, and the presence of this mutation in circulating VOCs led us to investigate the effect of a similar mutation on RaTG13. Using a set of sera (n=20) from HCWs who had been vaccinated with two doses of either BNT162b2 or AZD1222, we carried out pMN assays on SARS-CoV-2 and RaTG13 pseudotypes (WT and E484K or T484K, respectively). As expected, SARS-CoV-2 E484K neutralisation was significantly decreased when compared to SARS-CoV-2 (1.5 fold change, p=<0.0001) (Figure 2 G). Interestingly however, we observed the inverse trend with RaTG13 T484K, which showed a significant increase in neutralisation when compared to RaTG13 WT (2.0 fold change, p=<0.0014), highlighting that the significance of 484K changes to virus neutralisation are spike-background specific.

## Discussion

Within the last twenty years, three outbreaks of pathogenic coronaviruses have been recorded, with the current SARS-CoV-2 outbreak resulting in the global COVID-19 pandemic (17). Since the SARS-CoV outbreak in 2002-2003, significant focus has been placed on understanding the epidemiology of coronaviruses in order to assess risk and to prevent future outbreaks. This spurred large ecological surveillance studies, culminating in the discovery of numerous coronaviruses in bats, with one group reporting the detection of 293 coronaviruses from a single cave (3,18), nine of which were identified as betacoronaviruses, with one of these given the denominator RaTG13. Several bat coronaviruses, including RaTG13 were found to use the ACE2 receptor for entry and displayed broad species tropism (19–22). Since RaTG13 is the closest known relative of SARS-CoV-2 and binds to the human ACE2 receptor (14,22), our study sought to assess the level of cross-neutralisation of RaTG13 afforded by antibodies raised against SARS-CoV-2 either following natural infection, vaccination or both. Despite significant variation within the RBD we identified that SARS-CoV-2 specific immune sera generated through infection or immunisation is capable of neutralising RaTG13 pseudotypes. This has important implications for our understanding of betacoronavirus immunology and the future emergence of these viruses in humans, where herd immunity levels may be maintained at a high level.

Separately, within the current pandemic there are significant concerns that new variants of SARS-CoV-2 could arise that evolve to escape host-derived immunity. Our data shows the substantial reduction in neutralisation by the Beta VOC, B.1.351 (Figure 1C), which is consistent with others in the field (11,13,16,23) exemplifying the concern regarding immune escape. It is now well established that the E484K change present within the Beta RBD plays an important role in neutralising antibody evasion (24), data which we separately confirmed with a single point mutant (Figure 2G). Indeed, similar data exist for mutations at position N439 (15,25,26), Y449 (27), E484 (28), F486 (29), Q493 (30), N501 (31–33) and Y505 (34) within SARS-CoV-2. These sites also differ between the SARS-CoV-2 and RaTG13 RBD. With this in mind, it was therefore surprising to see that RaTG13 was efficiently neutralised by SARS-CoV-2 specific sera despite so many relevant changes in the RBD. These data indicate that SARS-CoV-2 immunity may tolerate significant changes to the RBD sequence without losing efficacy, and perhaps more optimistically that only a small number of changes exist which can dramatically alter neutralisation, conclusions which are supported by other recent observations (31).

There are various explanations which might explain why RaTG13 is more potently neutralised than SARS-CoV-2. A reduced receptor binding affinity for human ACE2 (5), when compared to SARS-CoV-2, might mean the RaTG13 Spike is more easily displaced from its receptor by competition from higher affinity antibodies. Supportive of this hypothesis, the N501D change in SARS-CoV-2, which we previously showed reduced human ACE2 receptor usage (7) increased neutralisation of this mutant (Figure 2F). The decreased neutralisation of the RaTG13 chimera which contains various substitutions that should enhance human ACE2 binding is also supportive of this “affinity model”. In contrast the T484K change in RaTG13, which we might expect to increase human ACE2 usage, actually increased neutralisation. However, it is likely the broader context for these changes in the overall structure of the RBD, and epitopes within, is also important. Data on SARS-CoV-1 and SARS-CoV-2 support a model that these viruses acquired higher human ACE2 usage during spill-over (7,35). If, as we suspect, lower affinity interactions are more sensitive to cross-neutralisation this could provide evidence for SARS-CoV-2 vaccination as a route to prevent subsequent spill-over of related sarbecoviruses. This is assuming that the majority of sarbecoviruses have an inherently lower affinity for the human ACE2 receptor whilst circulating in their natural reservoir.

In summary, our data shows that RaTG13 is potently neutralised by antibodies in convalescent sera from SARS-CoV-2 previously infected and/or vaccinated patients, suggesting that future potential spillover of RaTG13 or a closely related virus may be mitigated by pre-existing immunity to SARS-CoV-2 within the human population. How far this umbrella of cross-neutralisation extends across more distantly related sarbecoviruses is the source of continued research in our laboratory. Furthermore, the efficient neutralisation of RaTG13, despite its large number of RBD substitutions highlights that variation within the SARS-CoV-2 Spike may be, to a certain degree, controllable by existing vaccines and/or VOC-based boosters. It also provides evidence that the antigenically relevant amino acid mutations found in circulating variants, e.g. E484K, were generated as a result of antibody selection pressure and not genetic drift, since the time to the most recent common ancestor (TMRCA) of RaTG13 and SARS-CoV-2 is thought to be decades ago (36). Ultimately, our results suggest that the current priority should remain the effective identification and sequencing of SARS-CoV-2 variants, as it is these viruses which contain the most potent neutralisation-escape mutations.

## Acknowledgements

DB, JN, NT and AM were funded by The Pirbright Institute’s BBSRC institute strategic programme grant (BBS/E/I/00007038) and by the MRC funded grant G2P-UK; A National Virology Consortium to address phenotypic consequences of SARS-CoV-2 genomic variation, (MR/W005611/1). DC, MMN, AN, AC, PS, JCO, HB, NJT and JH are members of Humoral Immune Correlates to COVID-19 (HICC) consortium, funded by the UKRI and NIHR;(COV0170 - HICC: Humoral Immune Correlates for COVID19, Grant Reference code: MC_PC_20016). MMN and NJT are funded by Wellcome Trust/ UK FCDO (GB-CHC-210183). We thank the RPH Foundation Trust COVID-19 Research and Clinical teams, HCWs and Outpatients who participated in studies undertaken by the HICC.

## References

1. Zhou P, Yang X-L, Wang X-G, Hu B, Zhang L, Zhang W, et al. A pneumonia outbreak associated with a new coronavirus of probable bat origin. Nature. 2020 Mar;579(7798):270–3.

2. Zhu N, Zhang D, Wang W, Li X, Yang B, Song J, et al. A Novel Coronavirus from Patients with Pneumonia in China, 2019. New England Journal of Medicine. 2020 Feb 20;382(8):727–33.

3. Ge X-Y, Wang N, Zhang W, Hu B, Li B, Zhang Y-Z, et al. Coexistence of multiple coronaviruses in several bat colonies in an abandoned mineshaft. Virol Sin. 2016 Feb 18;31(1):31–40.

4. Lytras S, Hughes J, Martin D, Klerk A de, Lourens R, Pond SLK, et al. Exploring the natural origins of SARS-CoV-2 in the light of recombination. bioRxiv. 2021 May 27;2021.01.22.427830.

5. Wrobel AG, Benton DJ, Xu P, Roustan C, Martin SR, Rosenthal PB, et al. SARS-CoV-2 and bat RaTG13 spike glycoprotein structures inform on virus evolution and furin-cleavage effects. Nature Structural & Molecular Biology. 2020 Aug;27(8):763–7.

6. Walls AC, Park Y-J, Tortorici MA, Wall A, McGuire AT, Veesler D. Structure, Function, and Antigenicity of the SARS-CoV-2 Spike Glycoprotein. Cell. 2020 Apr 16;181(2):281-292.e6.

7. Conceicao C, Thakur N, Human S, Kelly JT, Logan L, Bialy D, et al. The SARS-CoV-2 Spike protein has a broad tropism for mammalian ACE2 proteins. PLOS Biology. 2020 Dec 21;18(12):e3001016.

8. Di Genova C, Sampson A, Scott S, Cantoni D, Mayora-Neto M, Bentley E, et al. Production, titration, neutralisation and storage of SARS-CoV-2 lentiviral pseudotypes. 2020 Dec 30 [cited 2021 Apr 25]; Available from: /articles/preprint/Production_titration_neutralisation_and_storage_of_SARS-CoV-2_lentiviral_pseudotypes/13502580/2

9. Ferrara F, Temperton N. Pseudotype Neutralization Assays: From Laboratory Bench to Data Analysis. Methods Protoc [Internet]. 2018 Jan 22 [cited 2021 Apr 25];1(1). Available from: https://www.ncbi.nlm.nih.gov/pmc/articles/PMC6526431/

10. Castillo-Olivares J, Wells D, Ferrari M, Chan A, Smith P, Nadesalingam A, et al. Towards Internationally standardised humoral Immune Correlates of Protection from SARS CoV 2 infection and COVID-19 disease. medRxiv. 2021 May 23;2021.05.21.21257572.

11. Zhou D, Dejnirattisai W, Supasa P, Liu C, Mentzer AJ, Ginn HM, et al. Evidence of escape of SARS-CoV-2 variant B.1.351 from natural and vaccine-induced sera. Cell. 2021 Apr 29;184(9):2348-2361.e6.

12. Cantoni D, Mayora-Neto M, Nadesalingham A, Wells DA, Carnell GW, Ohlendorf L, et al. Standardised, quantitative neutralisation responses to SARS-CoV-2 Variants of Concern by convalescent anti-sera from first wave infections of UK Health Care Workers and Patients. medRxiv. 2021 May 26;2021.05.24.21257729.

13. Hoffmann M, Arora P, Groß R, Seidel A, Hörnich BF, Hahn AS, et al. SARS-CoV-2 variants B.1.351 and P.1 escape from neutralizing antibodies. Cell [Internet]. 2021 Mar 20 [cited 2021 Apr 12]; Available from: https://www.sciencedirect.com/science/article/pii/S0092867421003676

14. Liu K, Pan X, Li L, Yu F, Zheng A, Du P, et al. Binding and molecular basis of the bat coronavirus RaTG13 virus to ACE2 in humans and other species. Cell. 2021 Jun 24;184(13):3438-3451.e10.

15. Thomson EC, Rosen LE, Shepherd JG, Spreafico R, Filipe A da S, Wojcechowskyj JA, et al. Circulating SARS-CoV-2 spike N439K variants maintain fitness while evading antibody-mediated immunity. Cell. 2021 Mar 4;184(5):1171-1187.e20.

16. Edara VV, Norwood C, Floyd K, Lai L, Davis-Gardner ME, Hudson WH, et al. Infection-and vaccine-induced antibody binding and neutralization of the B.1.351 SARS-CoV-2 variant. Cell Host & Microbe. 2021 Apr 14;29(4):516-521.e3.

17. Arora P, Jafferany M, Lotti T, Sadoughifar R, Goldust M. Learning from history: Coronavirus outbreaks in the past. Dermatologic Therapy. 2020;33(4):e13343.

18. Zhou P, Yang X-L, Wang X-G, Hu B, Zhang L, Zhang W, et al. Addendum: A pneumonia outbreak associated with a new coronavirus of probable bat origin. Nature. 2020 Dec;588(7836):E6–E6.

19. Ge X-Y, Li J-L, Yang X-L, Chmura AA, Zhu G, Epstein JH, et al. Isolation and characterization of a bat SARS-like coronavirus that uses the ACE2 receptor. Nature. 2013 Nov;503(7477):535–8.

20. Zheng M, Zhao X, Zheng S, Chen D, Du P, Li X, et al. Bat SARS-Like WIV1 coronavirus uses the ACE2 of multiple animal species as receptor and evades IFITM3 restriction via TMPRSS2 activation of membrane fusion. Emerging Microbes & Infections. 2020 Jan 1;9(1):1567–79.

21. Yang X-L, Hu B, Wang B, Wang M-N, Zhang Q, Zhang W, et al. Isolation and Characterization of a Novel Bat Coronavirus Closely Related to the Direct Progenitor of Severe Acute Respiratory Syndrome Coronavirus. Journal of Virology. 2016 Mar 15;90(6):3253–6.

22. Shang J, Ye G, Shi K, Wan Y, Luo C, Aihara H, et al. Structural basis of receptor recognition by SARS-CoV-2. Nature. 2020 May;581(7807):221–4.

23. Planas D, Bruel T, Grzelak L, Guivel-Benhassine F, Staropoli I, Porrot F, et al. Sensitivity of infectious SARS-CoV-2 B.1.1.7 and B.1.351 variants to neutralizing antibodies. Nature Medicine. 2021 Mar 26;1–8.

24. Wise J. Covid-19: The E484K mutation and the risks it poses. BMJ. 2021 Feb 5;372:n359.

25. Li Q, Wu J, Nie J, Zhang L, Hao H, Liu S, et al. The Impact of Mutations in SARS-CoV-2 Spike on Viral Infectivity and Antigenicity. Cell. 2020 Sep;182(5):1284-1294.e9.

26. Clark SA, Clark LE, Pan J, Coscia A, McKay LGA, Shankar S, et al. SARS-CoV-2 evolution in an immunocompromised host reveals shared neutralization escape mechanisms. Cell. 2021 May 13;184(10):2605-2617.e18.

27. Greaney AJ, Loes AN, Crawford KHD, Starr TN, Malone KD, Chu HY, et al. Comprehensive mapping of mutations in the SARS-CoV-2 receptor-binding domain that affect recognition by polyclonal human plasma antibodies. Cell Host & Microbe. 2021 Mar 10;29(3):463-476.e6.

28. Andreano E, Piccini G, Licastro D, Casalino L, Johnson NV, Paciello I, et al. SARS-CoV-2 escape in vitro from a highly neutralizing COVID-19 convalescent plasma. bioRxiv. 2020 Dec 28;2020.12.28.424451.

29. Liu Z, VanBlargan LA, Bloyet L-M, Rothlauf PW, Chen RE, Stumpf S, et al. Identification of SARS-CoV-2 spike mutations that attenuate monoclonal and serum antibody neutralization. Cell Host & Microbe. 2021 Mar 10;29(3):477-488.e4.

30. Starr TN, Greaney AJ, Addetia A, Hannon WW, Choudhary MC, Dingens AS, et al. Prospective mapping of viral mutations that escape antibodies used to treat COVID-19. Science. 2021 Feb 19;371(6531):850–4.

31. Rees-Spear C, Muir L, Griffith SA, Heaney J, Aldon Y, Snitselaar JL, et al. The effect of spike mutations on SARS-CoV-2 neutralization. Cell Reports. 2021 Mar 23;34(12):108890.

32. Wang Z, Schmidt F, Weisblum Y, Muecksch F, Barnes CO, Finkin S, et al. mRNA vaccine-elicited antibodies to SARS-CoV-2 and circulating variants. Nature. 2021 Feb 10;1–7.

33. Shen X, Tang H, McDanal C, Wagh K, Fischer W, Theiler J, et al. SARS-CoV-2 variant B.1.1.7 is susceptible to neutralizing antibodies elicited by ancestral spike vaccines. Cell Host & Microbe [Internet]. 2021 Mar 5 [cited 2021 Apr 12]; Available from: https://www.sciencedirect.com/science/article/pii/S1931312821001025

34. Rappazzo CG, Tse LV, Kaku CI, Wrapp D, Sakharkar M, Huang D, et al. Broad and potent activity against SARS-like viruses by an engineered human monoclonal antibody. Science. 2021 Feb 19;371(6531):823–9.

35. Wu K, Peng G, Wilken M, Geraghty RJ, Li F. Mechanisms of Host Receptor Adaptation by Severe Acute Respiratory Syndrome Coronavirus. J Biol Chem. 2012 Mar 16;287(12):8904–11.

36. Boni MF, Lemey P, Jiang X, Lam TT-Y, Perry BW, Castoe TA, et al. Evolutionary origins of the SARS-CoV-2 sarbecovirus lineage responsible for the COVID-19 pandemic. Nat Microbiol. 2020 Nov;5(11):1408–17.

